# p300/CBP-Driven Acetylation Stabilizes PARP7 to Reinforce Negative Feedback on Type I Interferon Signaling

**DOI:** 10.64898/2026.01.28.702217

**Authors:** Dan Li, Yao Liu, Ying Wang, Weijie Liu, Shihui Zhu, Ying Zhang, Luyi Wang, Xiaoqin Zhou, Wen Ding, Sanyong Zhu, Yongjun Dang, Jiaxue Wu

## Abstract

Immune suppression within the tumor microenvironment (TME) remains a major barrier to effective immunotherapy, yet the molecular mechanisms that constrain type I interferon (IFN-I) signaling in tumor cells are incompletely understood. PARP7 (also known as TIPARP), a mono-ADP-ribosyltransferase, has been reported to suppress IFN-I signaling by modifying TBK1. Here, we identify PARP7 as a previously unrecognized acetylation substrate of the histone acetyltransferases p300 and CBP. We show that p300/CBP-mediated acetylation of PARP7 at lysine 32 markedly enhances its stability, thereby reinforcing repression of the TBK1-IRF3-STAT1 axis and suppressing IFN-β-induced expression of interferon-stimulated genes, including CXCL10. Beyond their established function as transcriptional co-activators that support tumor cell proliferation, our findings uncover a direct role for p300/CBP in shaping innate antitumor immunity. Pharmacological inhibition of p300/CBP catalytic activity by A-485 abolishes PARP7 acetylation, accelerates its proteasomal turnover, and restores IFN-I signaling. In immunocompetent tumor models, this is accompanied by enhanced CD8^+^ T-cell infiltration and effector function. Together, these findings define a p300/CBP-PARP7 regulatory axis, uncover a non-canonical role for p300/CBP in innate immune suppression, and nominate this pathway as a tractable target for restoring antitumor immunity.

## Introduction

Current cancer immunotherapies have achieved durable clinical responses in a subset of patients; however, their overall efficacy is frequently limited by the immunosuppressive nature of the TME (de Visser & Joyce, 2023). Tumor-intrinsic mechanisms that attenuate innate immune sensing and prevent the initiation of productive antitumor immune responses therefore constitute a major barrier to therapeutic success (Pardoll, 2012). Among these, pathways controlling IFN-I production are of particular importance, as IFN-I signaling is essential for dendritic cell activation, antigen presentation, and the recruitment and functional licensing of cytotoxic T cells (Diamond *et al*, 2011; Montoya *et al*, 2002). Dissecting how IFN-I responses are actively restrained in tumor cells is thus critical for the rational development of strategies to overcome immune resistance.

The cGAS-STING pathway constitutes a central cytosolic DNA-sensing mechanism that links genomic instability to innate immune activation (Decout *et al*, 2021; Lu *et al*, 2026; Wang *et al*, 2025). Upon binding to cytosolic DNA, cGAS synthesizes the second messenger 2’3’-cGAMP, which activates STING and triggers the recruitment of TBK1. TBK1, in turn, phosphorylates IRF3 through the TBK1–IKKε complex and often engages parallel NF-κB signaling, culminating in the induction of IFN-β and a broad repertoire of interferon-stimulated genes (ISGs) (Ishikawa & Barber, 2008; Sun *et al*, 2013). These responses promote dendritic cell maturation, chemokine production, and T-cell recruitment, thereby facilitating productive antitumor immunity. In poorly immunogenic “cold” tumors, activation of the cGAS–STING axis can convert the TME into a more inflamed, “hot” state and improve responsiveness to immune checkpoint blockade (Girard *et al*, 2025).

Multiple tumor-intrinsic factors actively restrain cGAS-STING signaling, among which PARP7 has recently emerged as a key negative regulator (Gozgit *et al*, 2021; Popova *et al*, 2025). PARP7 is a member of the poly(ADP-ribose) polymerase family and functions primarily as a mono-ADP-ribosyltransferase, catalyzing NAD^+^-dependent mono-ADP-ribosylation of target proteins. It has been reported to modify TBK1, thereby establishing a negative feedback loop that suppresses IFN-I signaling activation (Sanderson *et al*, 2023; Yamada *et al*, 2016). Beyond this function, PARP7 has been implicated in the regulation of diverse transcription factors, including AHR, ERα, AR and HIF-1α, in a context-dependent manner (Jeltema *et al*, 2025; Zhang *et al*, 2020). Notably, PARP7 is an unusually short-lived protein, with reported half-lives on the order of minutes, resulting in extremely low steady-state abundance (Ikenga *et al*, 2025; Yang *et al*, 2023). This instability is reflected in the limited ability of currently available commercial antibodies to detect endogenous PARP7. Although PARP7 catalytic activity, including auto-modification, has been proposed to influence its turnover, the mechanisms governing PARP7 stability remain poorly understood.

In an effort to identify regulators of PARP7 stability, we focused on p300 and CBP, two closely related transcriptional co-activators with intrinsic histone acetyltransferase (HAT) activity. p300/CBP coordinate large-scale transcriptional programs by acetylating histones and numerous non-histone substrates, thereby shaping chromatin accessibility, cell identity, and stress responses (Goodman & Smolik, 2000; Liu *et al*, 2008; Martire *et al*, 2020). In cancer, the roles of p300/CBP are highly context-dependent: while they can act as tumor suppressors in certain settings, they frequently sustain oncogenic transcriptional programs and promote therapy resistance across diverse malignancies (Hogg *et al*, 2021). These observations have motivated the development of small-molecule p300/CBP inhibitors, including the catalytic HAT inhibitor A-485, which shows potent antitumor activity in preclinical models.

Beyond their tumor-intrinsic transcriptional functions, emerging evidence suggests that p300/CBP also influence tumor-immune interactions (Zeng *et al*, 2025); however, the molecular mechanisms by which they regulate innate immune signaling remain poorly defined. Here, we show that p300/CBP directly acetylate PARP7, thereby stabilizing the protein and reinforcing its inhibitory effect on the TBK1-IRF3-STAT1 axis. Conversely, pharmacological inhibition of p300/CBP by A-485 abolishes PARP7 acetylation, accelerates its proteasomal degradation, and restores IFN-I signaling.

Functionally, this leads to enhanced expression of IFN-β and downstream chemokines, such as CXCL10, and promotes CD8^+^ T-cell infiltration and effector activity in immunocompetent tumor models. Together, our findings uncover a previously unrecognized regulatory circuit linking p300/CBP to PARP7 and identify this axis as a key node controlling tumor-associated innate immune suppression.

## Results

### PARP7 directly interacts with p300/CBP

PARP7 functions as a negative regulator of IFN-I signaling in tumor cells by mono-ADP-ribosylating TBK1 and suppressing TBK1-IRF3-dependent transcription of IFNB1. Despite its established role as a central immunosuppressive factor, the molecular mechanisms governing PARP7 regulation remain incompletely defined. In particular, PARP7 is subject to tight post-translational control and undergoes rapid proteasomal turnover, raising the possibility that its stability and function are shaped by specific protein interactions. To delineate how PARP7 is dynamically regulated, we sought to define its protein interactome.

To identify proteins associated with PARP7, we expressed SBP-tagged PARP7 in HEK293T cells and performed streptavidin affinity purification followed by LC-MS/MS analysis. Gene ontology enrichment analysis revealed a significant overrepresentation of proteins involved in protein stability, transcriptional regulation, and chromatin-associated processes (Fig. 1A). Among the most highly enriched interactors were several E3 ubiquitin ligases, including HUWE1, UBR5, and RNF114 (Fig. 1B), suggesting that PARP7 turnover may be regulated through ubiquitin-proteasome pathways. Notably, the histone acetyltransferases p300 and CBP were also robustly enriched. Both proteins had previously been detected as PARP7-proximal in independent BioID datasets, providing additional confidence in the specificity of our proteomic results (Gibson *et al*, 2016). Consistent with this, genome-wide CRISPR resistance screens for the PARP7 inhibitor RBN-2397 identified *EP300* and *CREBBP* as significant hits, further implicating p300/CBP in PARP7 regulation (Chen *et al*, 2022).

**Figure 1.**
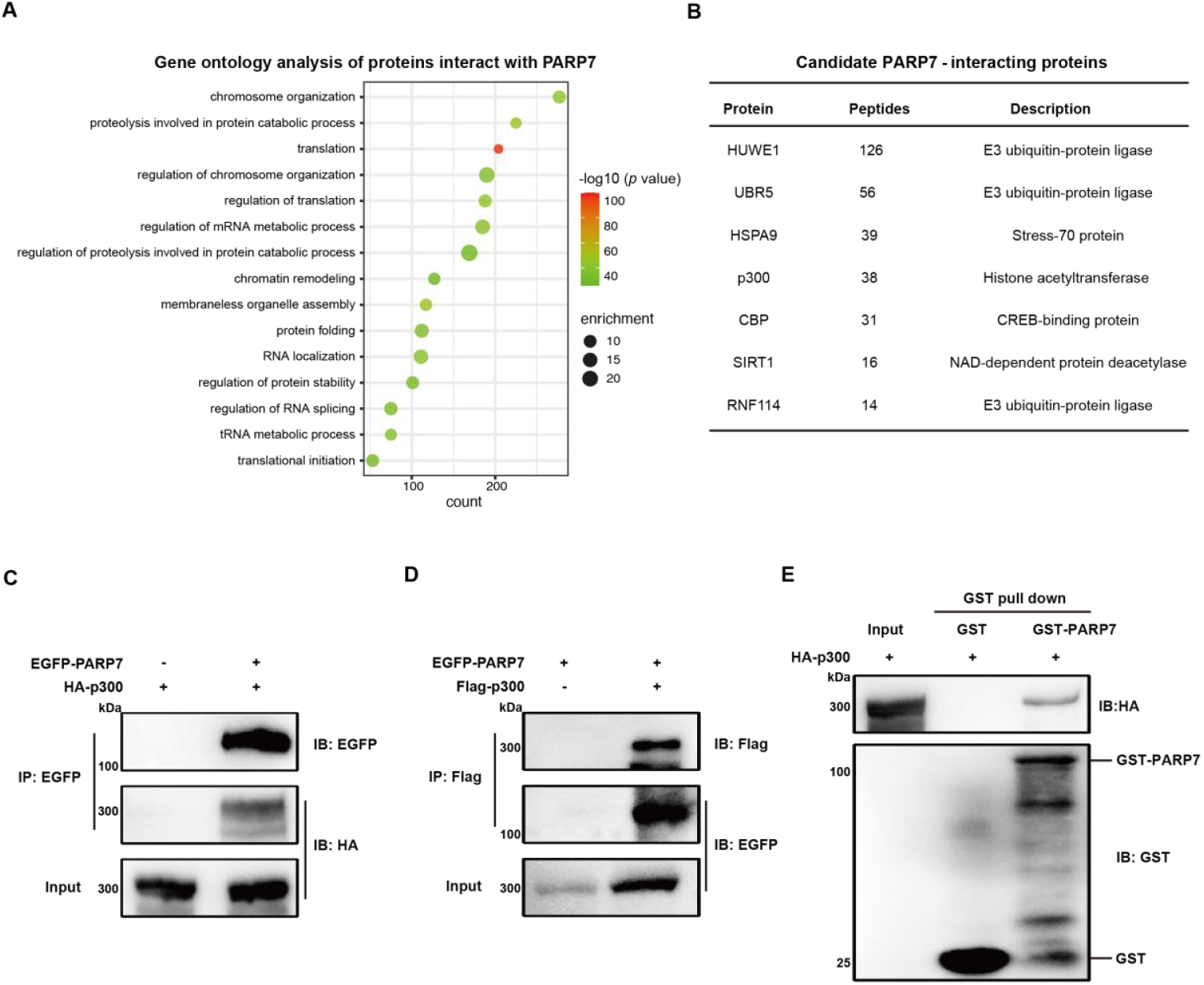
Identification and validation of p300/CBP as direct PARP7-interacting proteins. (A, B) SBP-tagged PARP7 was affinity-purified from HEK293T cells and analyzed by LC-MS/MS to determine associated proteins. Enriched pathways were assessed by gene ontology analysis (A), and representative high-confidence candidates are listed (B). Bubble size indicates enrichment magnitude, and color represents -log10 (p value). (C, D) The interaction between PARP7 and p300/CBP was validated by reciprocal co-immunoprecipitation in HEK293T cells. Co-immunoprecipitation of EGFP-PARP7 with HA-p300 (C) and reverse co-immunoprecipitation of Flag-p300 with EGFP-PARP7 (D). (E) In vitro GST pull-down using purified GST-PARP7 and HA-p300. GST-PARP7 was expressed in *E*.*coli*, whereas HA-p300 was expressed in HEK293T cells. All Western blot data shown are representative of three independent experiments.

We next validated the interaction between PARP7 and p300/CBP by co-immunoprecipitation. In HEK293T cells co-expressing EGFP-PARP7 and either HA-p300 or Flag-p300, immunoprecipitation of EGFP-PARP7 efficiently recovered p300, and reciprocal pull-down of p300 enriched PARP7 (Fig. 1C,D). To determine whether this interaction is direct, we performed in vitro pull-down assays using purified GST-PARP7, which demonstrated direct binding to HA-p300 (Fig. 1E). Together, these data establish p300/CBP as bona fide direct interaction partners of PARP7.

A recent preprint independently reported that PARP7 mono-ADP-ribosylates p300 and CBP, identifying these acetyltransferases as direct functional targets of PARP7 (Siordia *et al*, 2025). Consistent with this report, our PARP7 interactome was enriched for p300/CBP, as well as SIRT1, a NAD^+^-dependent deacetylase. The presence of enzymes that both install and remove lysine acetylation suggested that PARP7 itself may be subject to acetylation and dynamic post-translational regulation, a possibility we explored in subsequent experiments.

### PARP7 is an acetylation substrate of p300/CBP

Given the direct interaction between PARP7 and p300/CBP, we next examined whether PARP7 is acetylated in cells. To increase detection sensitivity, we generated HEK293T cells stably expressing SBP-tagged PARP7, and enriched the protein by streptavidin affinity purification prior to immunoblotting with a pan-acetyl-lysine antibody. PARP7 displayed detectable basal acetylation under steady-state conditions (Fig. 2A). Ectopic expression of p300 markedly increased PARP7 acetylation in a dose-dependent manner, whereas a catalytically inactive p300 mutant (Myc-p300 S1396R/Y1397R) failed to do so (Fig. 2B, 2C), indicating that p300 acetyltransferase activity is required for this modification.

**Figure 2.**
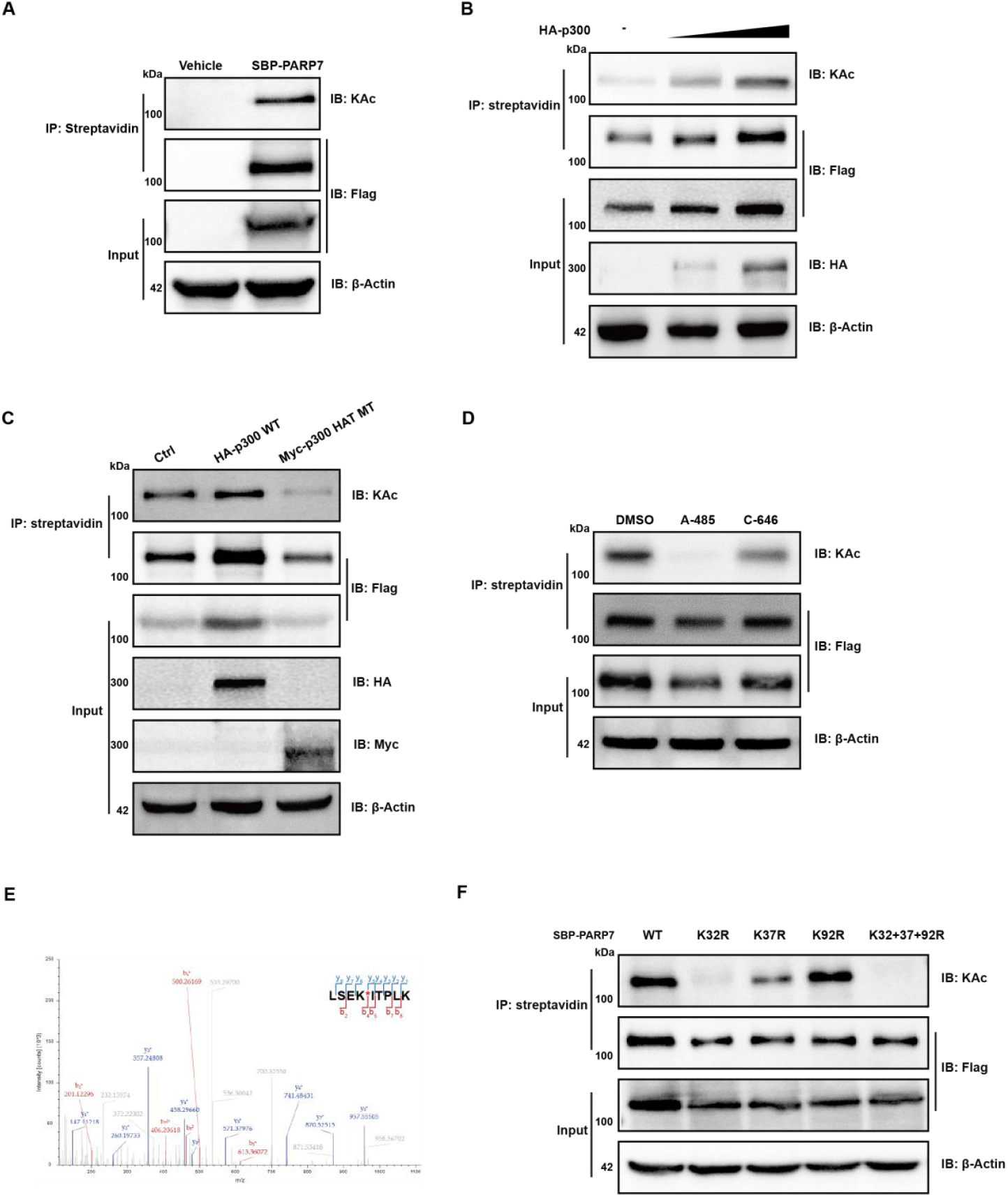
PARP7 serves as an acetylation substrate of p300/CBP. (A) Streptavidin affinity enrichment of SBP-PARP7 from HEK293T cells stably expressing SBP-PARP7, followed by immunoblotting to assess its lysine acetylation. Endogenous PARP7 acetylation was detectable without exogenous p300. (B) Increasing amounts of HA-p300 were expressed in HEK293T cells stably expressing SBP-PARP7. Streptavidin pulldown showed dose-dependent enhancement of PARP7 acetylation. (C) Wild-type HA-p300 but not the catalytic mutant Myc-p300 (S1396R/Y1397R) increased PARP7 acetylation, demonstrating dependence on p300 HAT activity. SBP-PARP7-expressing HEK293T cells were transfected with HA-p300 wild-type or Myc-p300 HAT-mutant (S1396R Y1397R) constructs. Streptavidin pull-down was performed prior to immunoblot analysis. (D) Treatment with the selective p300/CBP inhibitor A-485 nearly abolished PARP7 acetylation, whereas C-646 produced a weaker reduction. SBP-PARP7-expressing HEK293T cells were treated with DMSO, 1 μM A-485 or 10 μM C-646 for 24 hours, followed by streptavidin pull-down and immunoblot detection of PARP7 acetylation. (E) LC-MS/MS identification of PARP7 lysine 32 acetylation. Representative mass spectrometry spectrum showing the acetylated K32 peptide from purified PARP7. Purified SBP-PARP7 peptides were analyzed by tandem mass spectrometry, and acetylated residues were assigned based on MS/MS fragmentation spectra. (F) Mutation of individual lysines to arginine revealed that K32R strongly reduces PARP7 acetylation. SBP-PARP7 wild-type or lysine-to-arginine mutants (K32R, K37R, K92R, or combined K32+37+92R) were expressed in HEK293T cells. Following streptavidin pull-down, acetylation levels of the purified proteins were examined by immunoblotting. All Western blot data shown are representative of three independent experiments.

Because p300 and CBP share extensive structural homology and largely overlapping substrate specificities, we treated them as a functional unit and asked whether PARP7 acetylation depends on their shared catalytic activity (Jin *et al*, 2011; Liu *et al*., 2008; Wang *et al*, 2013). Pharmacological inhibition with the selective p300/CBP catalytic inhibitor A-485 nearly abolished PARP7 acetylation, whereas the less potent inhibitor C-646 caused only a partial reduction (Fig. 2D). Consistent with contributions from both enzymes, genetic deletion of EP300 reduced—but did not eliminate—PARP7 acetylation (Fig. S1A,B), while A-485 treatment produced a more pronounced effect. Together, these data indicate that PARP7 acetylation is redundantly mediated by p300 and CBP.

To identify the acetylation sites, we performed LC-MS/MS analysis of purified PARP7 and identified several candidate acetylated lysines. Three high-confidence sites—K32, K37, and K92—were selected for further validation (Fig. 2E; Fig. S1C). Each site was individually mutated to arginine (K to R) to mimic a non-acetylatable state. The K32R mutation markedly reduced the overall acetylation level of PARP7, whereas K37R and K92R exerted only mild effects (Fig. 2F). These data identify K32 as the dominant p300/CBP-dependent acetylation site on PARP7.

Together, these findings demonstrate that PARP7 forms a stable complex with p300/CBP and is directly acetylated by their shared catalytic activity, with K32 serving as the principal acetylation site.

### p300/CBP‐mediated acetylation enhances PARP7 protein stability

Building on these observations, reduced PARP7 acetylation was consistently accompanied by a decrease in PARP7 protein abundance, whether achieved by pharmacological inhibition with A-485 or by mutation of the major acetylation site K32. This convergence prompted us to examine whether p300/CBP regulates PARP7 stability in an acetyltransferase-dependent manner. Inhibition of p300/CBP catalytic activity with A-485 consistently reduced PARP7 protein abundance, without affecting PARP7 mRNA levels (Fig. 3A; Fig. S2A), suggesting post-translational regulation. Conversely, ectopic expression of p300 increased PARP7 protein levels in a dose-dependent manner (Fig. 3B). To determine whether p300/CBP-mediated acetylation protects PARP7 from proteasomal degradation, we treated cells with the proteasome inhibitor MG132. MG132 stabilized PARP7 under basal conditions and largely prevented the loss of PARP7 induced by A-485 (Fig. 3C), indicating that acetylation limits proteasome-dependent turnover.

**Figure 3.**
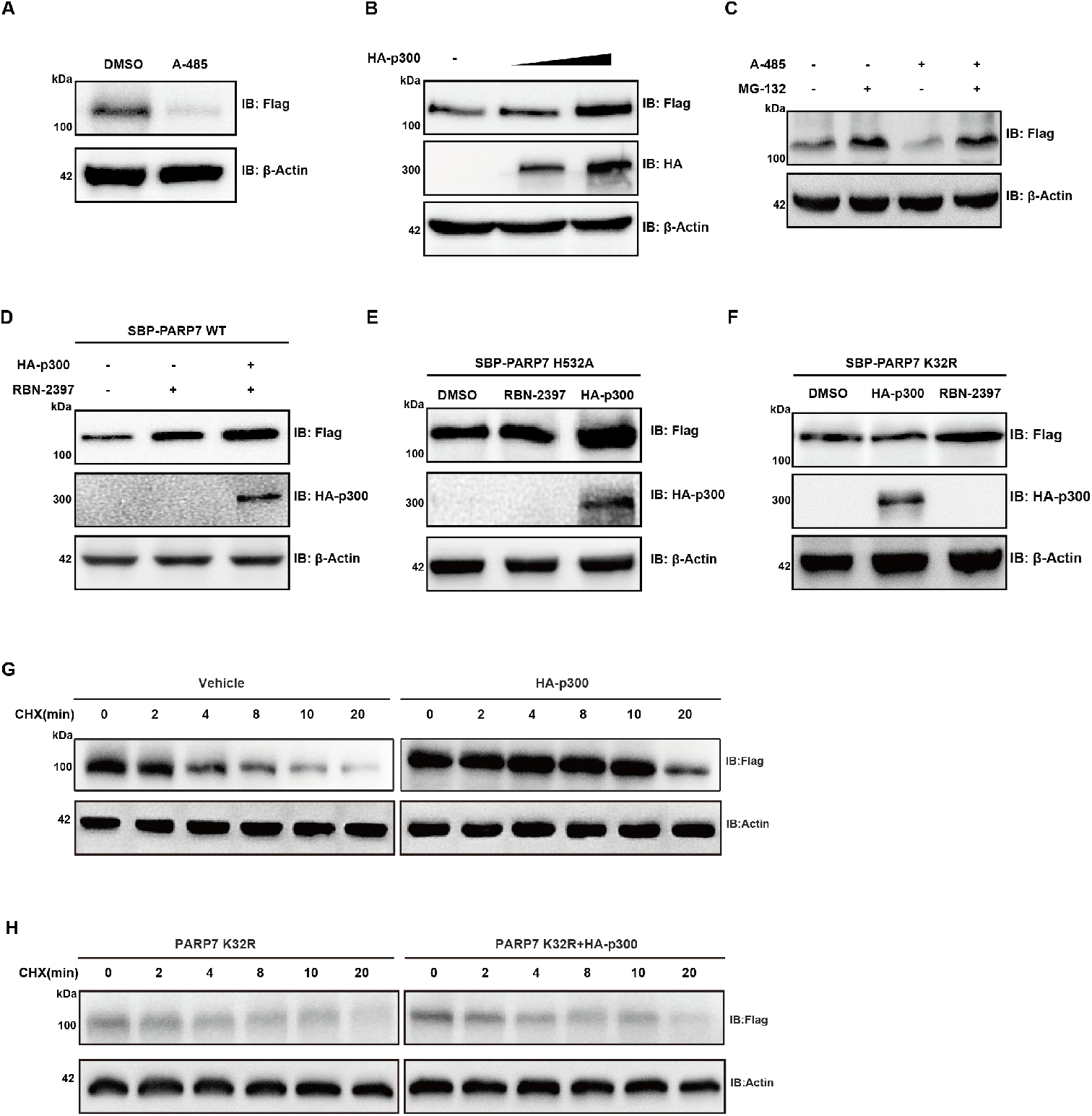
p300/CBP-mediated acetylation stabilizes PARP7. (A)Cells treated with A-485 (1 μM, 24 h) exhibit a marked reduction in PARP7 protein compared with DMSO control, indicating that p300/CBP catalytic activity is required to maintain PARP7 abundance. (B) HEK293T cells stably expressing SBP-PARP7 were transfected with increasing amounts of HA-p300. Immunoblotting of whole-cell lysates shows that PARP7 protein levels increase in a dose-dependent manner upon HA-p300 overexpression. (C) Proteasome inhibition rescues PARP7 loss. MG132 (5 μM, 6 h) stabilizes PARP7 under basal conditions and nearly abolishes the A-485–induced reduction, demonstrating that loss of acetylation accelerates proteasomal degradation. (D) Overexpression of HA-p300 enhances the RBN-2397-induced accumulation of PARP7 protein, as shown by immunoblotting of whole-cell lysates from HEK293T cells stably expressing SBP-PARP7. (E) The catalytically inactive PARP7 H532A mutant, when transiently expressed in HEK293T cells, no longer responds to RBN-2397 treatment, whereas its protein abundance is still increased upon HA-p300 overexpression. This result separates the stabilizing effect of p300-mediated acetylation from the auto-MARylation-dependent stabilization. (F) In the non-acetylatable K32R mutant, p300 fails to stabilize PARP7, while RBN-2397 still increases protein levels, confirming that K32 acetylation is essential for p300-dependent stabilization but dispensable for MARylation-mediated effects. (G-H) Cycloheximide chase assay showing that PARP7 has a short half-life under basal conditions but is markedly stabilized by p300 overexpression (G). In contrast, this stabilizing effect was lost on PARP7 K32R (H). PARP7 decay was monitored by immunoblotting across time points. All Western blot data shown are representative of three independent experiments.

PARP7 has been reported to undergo auto-mono-ADP-ribosylation, which contributes to its intrinsic instability. To assess how this mechanism intersects with acetylation, we examined the effects of the PARP7 catalytic inhibitor RBN-2397 and the catalytically inactive PARP7 mutant (H532A). In cells expressing wild-type PARP7, RBN-2397 increased PARP7 protein abundance, and p300 overexpression further enhanced this effect (Fig. 3D). In contrast, RBN-2397 failed to stabilize the H532A mutant, whereas p300 overexpression remained effective (Fig. 3E), indicating that p300-mediated stabilization occurs independently of PARP7 catalytic activity. Conversely, mutation of the acetylation site (K32R) abolished the stabilizing effect of p300 while preserving responsiveness to RBN-2397 (Fig. 3F). These observations together demonstrate that p300/CBP-mediated acetylation and PARP7 auto-MARylation independently shape PARP7 protein stability. This conclusion further supported by the finding that p300 overexpression does not interfere with PARP7 auto-ADP-ribosylation (Fig. S2B).

To directly measure the impact of acetylation on PARP7 turnover, we performed cycloheximide chase assays. Under control conditions, PARP7 exhibited a short half-life of approximately four minutes, whereas p300 overexpression markedly prolonged its half-life to nearly twenty minutes (Fig. 3G). In contrast, this stabilizing effect was lost upon mutation of the major acetylation site K32, as p300 overexpression failed to extend the half-life of the PARP7 K32R mutant (Fig. 3H). Thus, p300/CBP-mediated acetylation stabilizes PARP7 by protecting it from rapid proteasomal degradation.

Collectively, these results establish that p300/CBP acetylates PARP7—primarily at K32—to enhance its stability by preventing proteasome-dependent degradation. This mechanism provides a molecular explanation for how p300/CBP contributes to sustaining PARP7’s suppressive effect on type I interferon signaling.

### p300/CBP enhance the suppression of the IFN‐I pathway by stabilizing PARP7

We next examined the functional consequences of p300/CBP-mediated PARP7 stabilization. Given the established role of PARP7 as a negative regulator of IFN-I signaling through mono-ADP-ribosylation of TBK1, we asked whether inhibition of p300/CBP would phenocopy PARP7 inhibition at the level of TBK1 regulation. To this end, we assessed TBK1 ADP-ribosylation in CT26, MB49, and PC3 cells after A-485 treatment (1 μM, 24 h). ADP-ribosylated proteins were enriched using GST-Macro(PARP14 MacroD2) pull-down assays, which selectively capture mono-ADP-ribosylated substrates. In all three cell lines, pharmacological inhibition of PARP7 with RBN-2397 markedly reduced TBK1 ADP-ribosylation, and A-485 produced a comparably strong decrease (Fig. 4A). Loss of TBK1 ADP-ribosylation was accompanied by a robust increase in TBK1 phosphorylation (Fig. 4B), indicating relief of PARP7-mediated repression.

**Figure 4.**
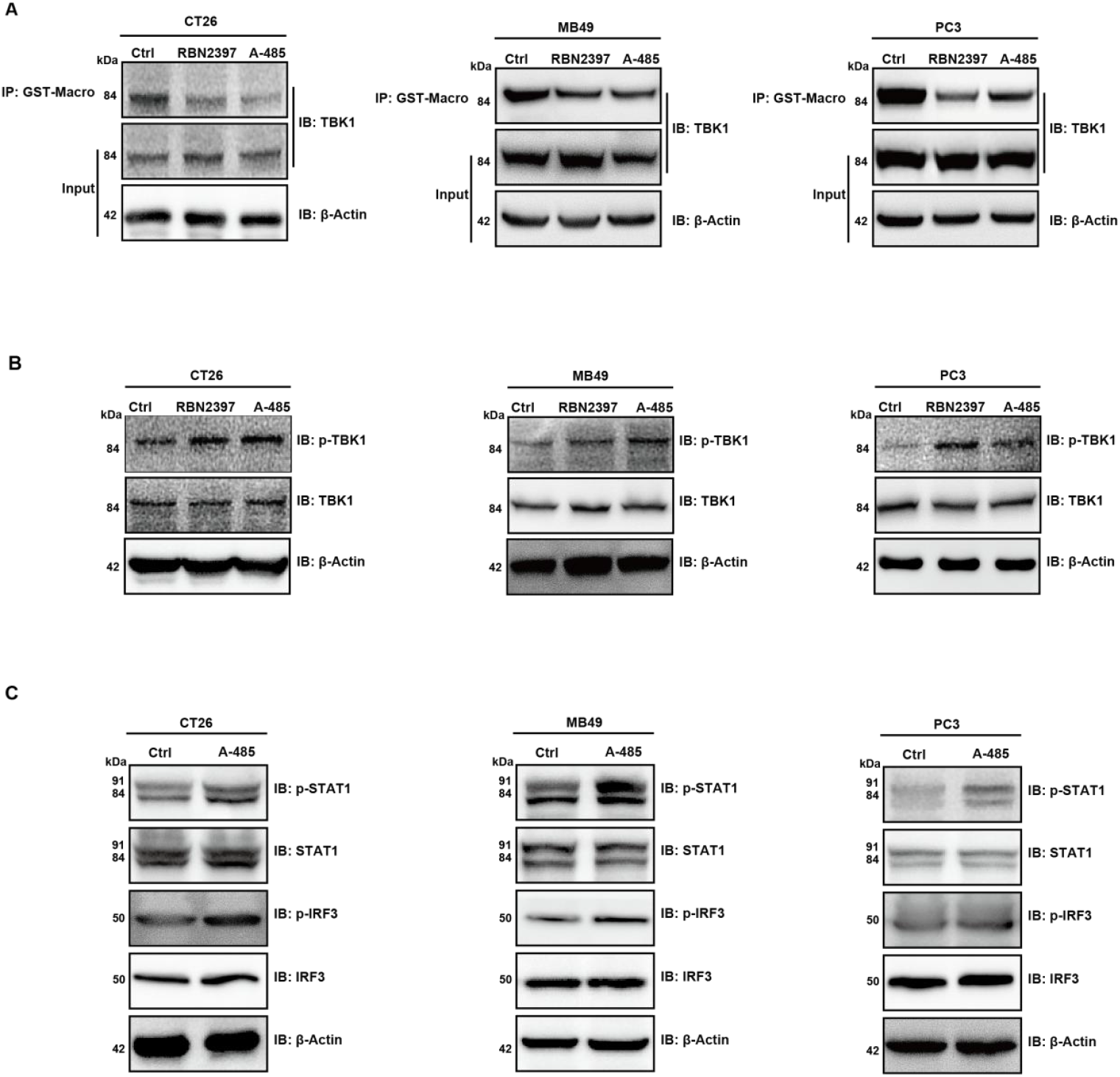
p300/CBP stabilize PARP7 to enhance suppression of IFN-I signaling. (A) A-485 or RBN-2397 reduces TBK1 ADP-ribosylation. CT26, MB49, and PC3 cells were treated with 0.5 μM RBN-2397 or 1 μM A-485 for 24h. ADP-ribosylated proteins were enriched using GST-Macro pull-down and analyzed by immunoblotting for TBK1. (B) Inhibition of p300/CBP enhances TBK1 phosphorylation. Lysates from CT26, MB49, and PC3 cells treated as in (A) were analyzed by immunoblotting for phosphorylated TBK1 (p-TBK1). (C) A-485 activates downstream IFN-I signaling components IRF3 and STAT1. A-485 increased phosphorylation of IRF3 and STAT1 in CT26 and MB49 cells; PC3 cells showed a weaker but detectable response. All Western blot data shown are representative of three independent experiments.

To further validate the impact of A-485 on downstream components of the IFN-I pathway, we examined IRF3 and STAT1 phosphorylation. A-485 treatment (1 μM, 24 h) increased phosphorylation of both IRF3 and STAT1 in CT26 and MB49 cells (Fig. 4C), confirming effective activation of the IFN-I signaling cascade. Although PC3 cells are known to exhibit attenuated IFN-I responsiveness and reduced sensitivity to PARP7 inhibition, A-485 nonetheless induced detectable increases in IRF3 and STAT1 phosphorylation, albeit to a lesser extent (Fig. 4C).

To determine whether these effects require PARP7, we generated *PARP7*-knockout MB49 cells (Fig. S3A). In the absence of PARP7, A-485 failed to enhance STAT1 phosphorylation (Fig. S3B), demonstrating that the effects of p300/CBP inhibition on IFN-I signaling require PARP7 expression.

Together, these data indicate that p300/CBP suppress IFN-I signaling by stabilizing PARP7. Inhibition of p300/CBP catalytic activity destabilizes PARP7, reduces TBK1 ADP-ribosylation, and restores TBK1-IRF3-STAT1 signaling, thereby relieving PARP7-dependent repression of the IFN-I pathway.

### Inhibition of p300/CBP promotes interferon‐stimulated genes expression

Having established that p300/CBP inhibition destabilizes PARP7 and reactivates proximal IFN-I signaling, we next asked whether this translated into enhanced transcription of interferon-stimulated genes (ISGs). Across multiple tumor cell lines—including CT26, MB49, ID8, and PC3—treatment with A-485 consistently increased mRNA levels of IFNB and the IFN-inducible chemokine CXCL10, indicating broad activation of the IFN-I transcriptional program. In CT26 cells, induction of IFNB and CXCL10 by A-485 was less pronounced than that achieved with the PARP7 inhibitor RBN-2397 (Fig. S4A). In contrast, in MB49, ID8, and PC3 cells, A-485 elicited substantially stronger upregulation of IFNB and CXCL10 than RBN-2397 (Fig. 5A, 5B). This difference was particularly evident in PC3 cells, which have previously been reported to respond poorly to RBN-2397 unless PARP7 expression is experimentally enhanced. These observations suggest that, in certain cellular contexts, reducing PARP7 protein abundance is more effective at unleashing IFN-I-driven transcription than inhibiting PARP7 catalytic activity alone.

**Figure 5.**
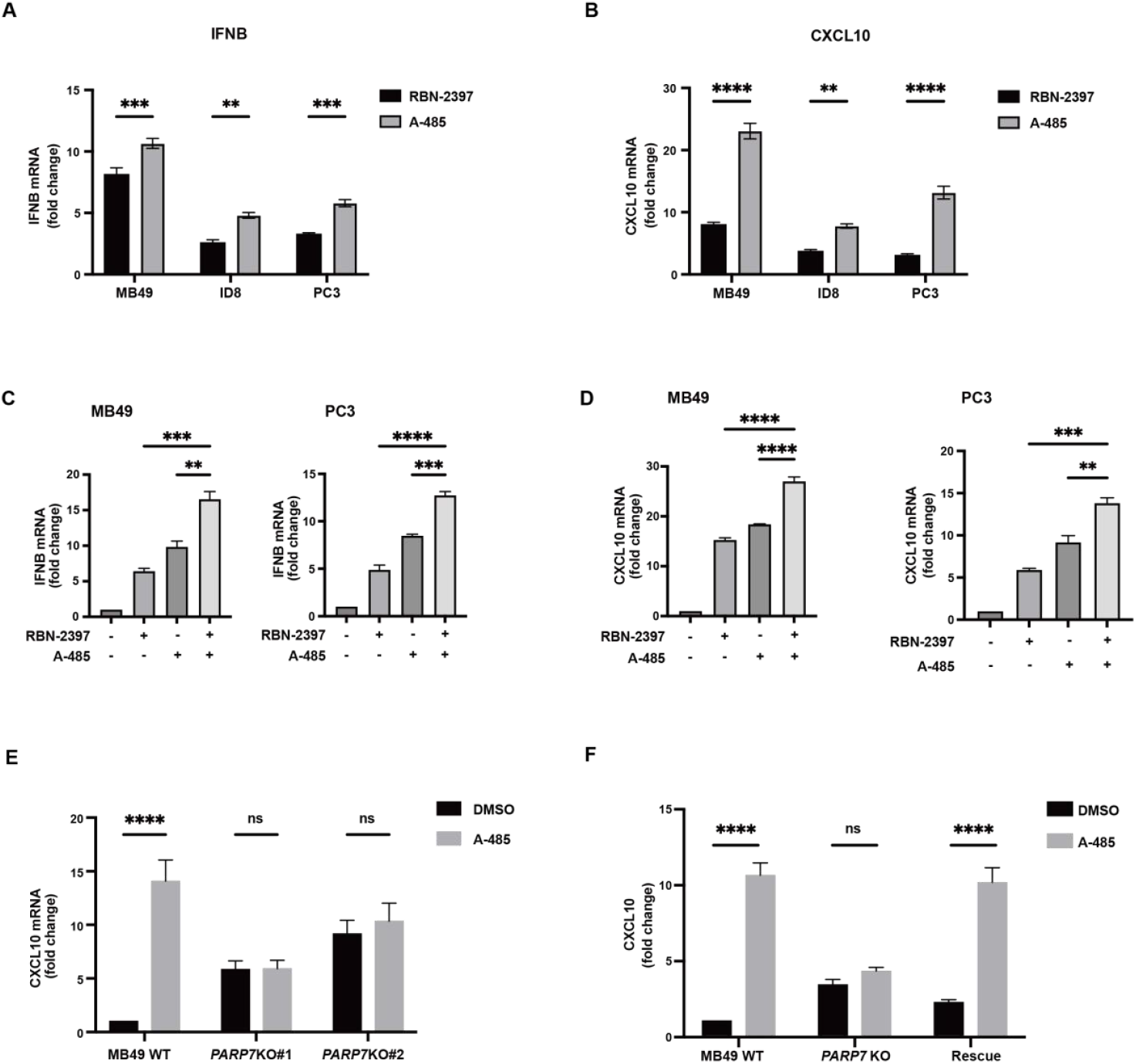
Inhibition of p300/CBP enhances IFN-I signaling and synergizes with RBN-2397. (A, B) qRT–PCR analysis of IFNB (A) and CXCL10 (B) mRNA expression in MB49, ID8, and PC3 cells following treatment with RBN-2397 (0.5 μM, 24h) or A-485 (1 μM, 24h). A-485 induced stronger IFN-I-responsive gene expression than RBN-2397 in all three cell lines. (C–D) qRT–PCR analysis of IFNB (C) and CXCL10 (D) mRNA levels in MB49 and PC3 cells treated with RBN-2397 (0.5 μM, 24h), A-485 (1 μM, 24h), or the combination. Combined inhibition of p300/CBP and PARP7 resulted in markedly greater induction of IFN-I-responsive genes than either treatment alone, indicating functional synergy. (E) qRT–PCR analysis of CXCL10 mRNA expression in MB49 wild-type and *PARP7*-knockout cells treated with DMSO or A-485 (1 μM, 24h). Loss of PARP7 substantially attenuated A-485-mediated induction of CXCL10. (F) qRT–PCR analysis of CXCL10 mRNA levels in MB49 wild-type, *PARP7*-knockout, and PARP7-rescued cells treated with DMSO or A-485 (1 μM, 24h). Re-expression of wild-type PARP7 restored responsiveness to A-485, demonstrating that A-485–induced CXCL10 expression depends on PARP7. Data shown are from one representative experiment (mean ± SEM of n = 3 technical replicates), and similar results were obtained in two additional independent experiments. Statistical significance was determined by one-way ANOVA. ns, not significant; *p < 0.05, **p < 0.01, ***p < 0.001, ****p < 0.0001

We next examined whether simultaneous inhibition of p300/CBP and PARP7 produced additive effects. In MB49 and PC3 cells, which responded most robustly to A-485, combined treatment with A-485 and RBN-2397 resulted in markedly greater induction of IFNB and CXCL10 than either agent alone (Fig. 5C, 5D). A similar cooperative effect was also observed in CT26 and ID8 cells (Fig. S4B, S4C). These suggests potential functional synergy between p300/CBP inhibition and PARP7 inhibition.

To determine whether A-485-induced ISG expression requires PARP7, we treated *PARP7*-knockout cells with A-485. Loss of PARP7 substantially blunted the ability of A-485 to induce CXCL10 expression (Fig. 5E). Reintroduction of PARP7 into these PARP7-deficient cells largely restored responsiveness to A-485 (Fig. 5F), confirming that the effects of A-485 depend on the presence of PARP7 protein.

Together, these findings demonstrate that p300/CBP activity sustains PARP7 protein stability and reinforces PARP7-mediated negative feedback on the IFN-I pathway. Inhibition of p300/CBP destabilizes PARP7, releasing its brake on IFN-I signaling and enabling stronger expression of interferon-stimulated genes (ISGs), including CXCL10.

### In vivo p300/CBP inhibition enhances CD8^+^ T cell‐mediated antitumor immunity

To determine the relevance of the p300/CBP-PARP7 axis in regulating antitumor immunity in vivo, we established a subcutaneous MB49 tumor model in C57BL/6 mice and performed a systematic pharmacological intervention. MB49 cells (1×10^6^ in 100 μL PBS) were injected subcutaneously into the right flank of mice, and tumor growth was monitored daily. Once most tumors reached a stable volume of approximately 80 mm^3^, mice were randomized into control and A-485 treatment groups based on tumor size to ensure comparable baselines (Fig. 6A). Daily administration of A-485 (20 mg/kg, intraperitoneally) significantly suppressed tumor growth compared with controls (Fig. 6B). Tumor growth curves showed that A-485 markedly slowed progression, with significant divergence from day 3 post-treatment (Fig. 6C).

**Figure 6.**
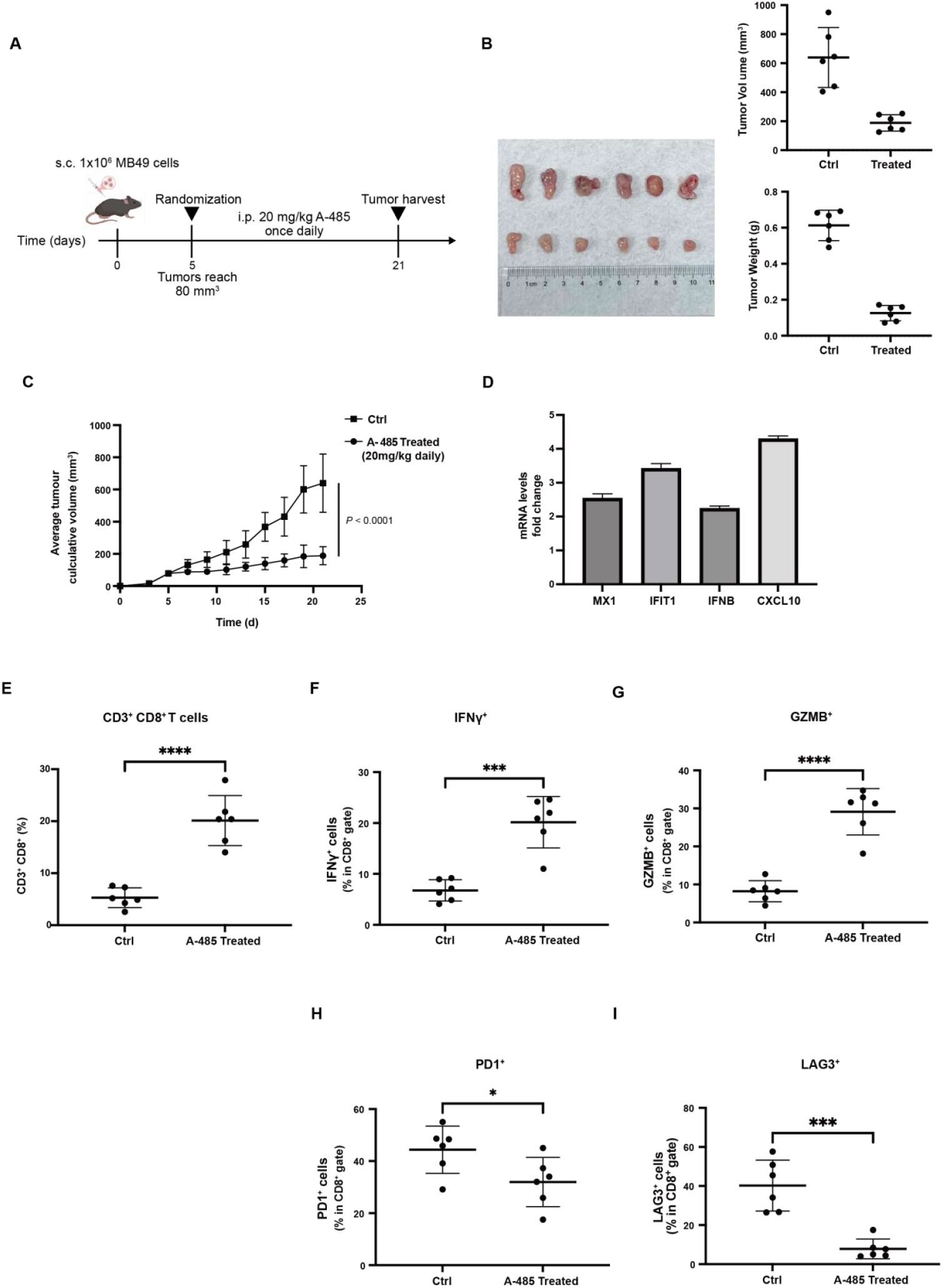
In vivo p300/CBP inhibition enhances CD8^+^ T cell-mediated antitumor immunity. (A) Experimental scheme of MB49 subcutaneous tumor model and A-485 treatment. MB49 cells (1×10^6^) were injected subcutaneously into the right flank of C57BL/6 mice (8 weeks), followed by randomization when tumors reached ∼80 mm^3^ and daily intraperitoneal administration of A-485 (20 mg/kg). Tumors were harvested at the study endpoint. (B) Representative images of excised tumors (left) and quantification of tumor volume and weight (right) from control and A-485-treated mice at endpoint, showing a marked reduction in tumor burden upon A-485 treatment. Data represent mean ± SD (n = 6 mice per group). (C) Tumor growth curves over time. Tumor volumes were monitored every two days. A-485 treatment markedly slowed tumor progression, with significant divergence observed from day 3 post-treatment. Data are presented as mean ± SEM (n = 6 mice per group). Statistical significance was determined by two-way ANOVA with repeated measures. (D) qRT–PCR analysis of IFN-I–responsive genes (MX1, IFIT1, IFNB, and CXCL10) in tumor tissues. Data shown are from one representative experiment (mean ± SEM of n = 3 technical replicates), and similar results were obtained in another independent experiment. (E–G) A-485 enhances CD8^+^ T cell infiltration and effector function. Tumor-infiltrating lymphocytes were isolated and analyzed by flow cytometry. (E) Proportion of CD3^+^CD8^+^T cells; (F) Frequency of IFN-γ^+^ and (G) GZMB^+^ CD8^+^ T cells. (H-I) A-485 reduces CD8^+^ T cell exhaustion markers. Expression of PD-1 (H) and LAG-3 (I) on tumor-infiltrating CD8^+^ T cells was assessed by flow cytometry, showing decreased exhaustion in A-485-treated mice. Data in (E-I) are presented as mean ± SEM (n = 6 mice per group). Statistical significance was determined by unpaired two-tailed Student’s t-test. ns, not significant; *p < 0.05, **p < 0.01, ***p < 0.001, ****p < 0.0001

To investigate whether the antitumor effect was linked to enhanced immune responses, tumors were first harvested and subjected to qPCR analysis. A-485 treatment resulted in a 2 to 4-fold upregulation of multiple IFN-I-responsive genes, including MX1, IFIT1, IFNB, and CXCL10 (Fig. 6D), indicating activation of intratumoral IFN-I signaling.

To determine whether this transcriptional reprogramming translated into altered immune cell infiltration, tumors were subsequently processed by Percoll density gradient centrifugation to isolate tumor-infiltrating leukocytes, followed by multiparameter flow cytometric analysis. Consistent with the elevated expression of IFN-I-induced chemokines, A-485 treatment significantly increased the proportion of infiltrating CD3^+^CD8^+^ T cells compared with control tumors (Fig. 6E). Moreover, CD8^+^ T cells from A-485-treated tumors displayed enhanced effector function, as evidenced by increased frequencies of IFN-γ^+^ and GZMB^+^cells (Fig. 6F, 6G), accompanied by reduced expression of exhaustion markers PD-1 and LAG-3 (Fig. 6H, 6I).

Collectively, our in vivo data demonstrate that the p300/CBP inhibitor A-485 reshapes the tumor immune microenvironment by activating IFN-I related pathways, enhancing CD8^+^ T cell infiltration and effector function while reducing exhaustion phenotypes. This coordinated immune reprogramming effectively suppresses MB49 tumor growth and provides compelling evidence that the p300/CBP-PARP7 axis plays a critical role in tumor immune evasion.

## Discussion

In this study, we uncover a novel mechanism linking the lysine acetyltransferases p300/CBP with innate immune suppression in tumor cells. We show that p300/CBP directly acetylate PARP7 (at K32), markedly delaying its proteasomal degradation and thereby elevating its steady-state levels. Because PARP7 mono-ADP-ribosylates TBK1 (Yamada *et al*., 2016) and inhibits its activation of IRF3-STAT1(Jeltema *et al*., 2025), this p300/CBP-mediated acetylation of PARP7 effectively enforces a brake on the TBK1-IRF3-STAT1 axis and restrains IFN-β production (Fig. 7). To our knowledge, this is the first demonstration that PARP7 is a direct substrate of p300/CBP. Importantly, these findings extend p300/CBP’s repertoire beyond classical transcriptional co-activation: they reveal a direct post-translational route by which p300/CBP can dampen antiviral-type I IFN signaling (Jin *et al*, 2017).

**Figure 7.**
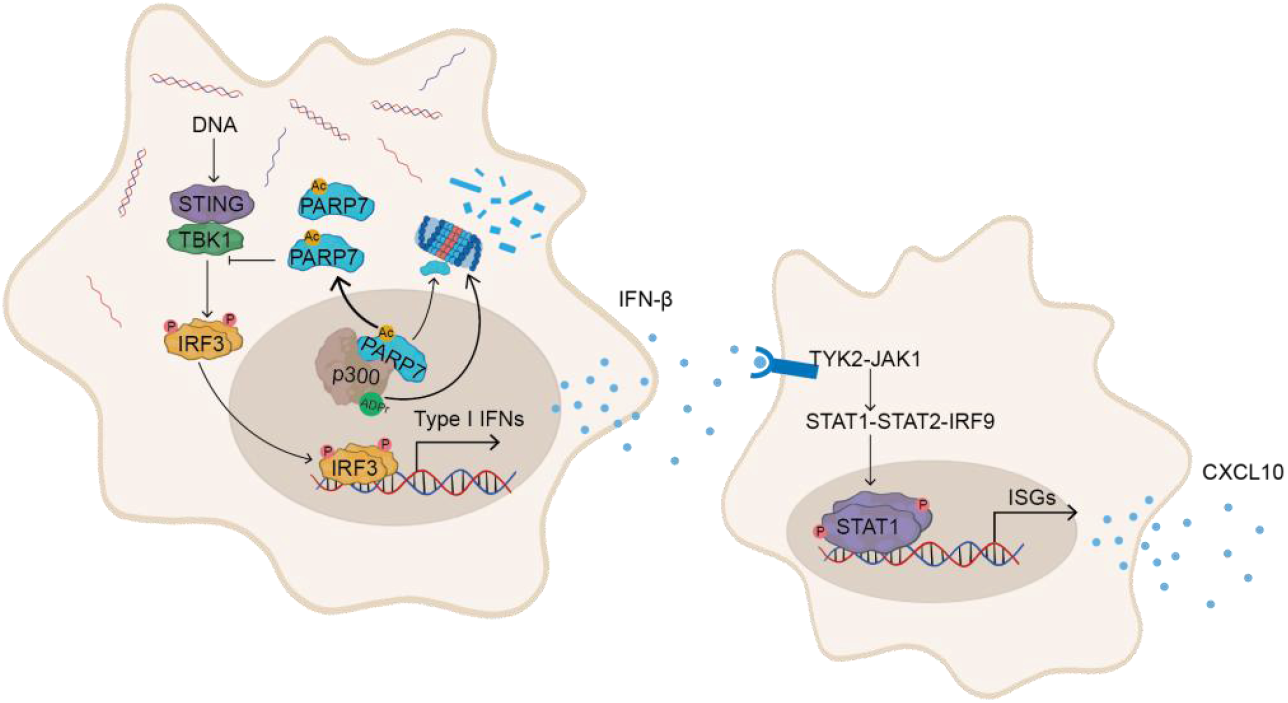
Model of p300/CBP-PARP7 axis. p300/CBP directly acetylate PARP7 at lysine 32, thereby stabilizing the protein and reinforcing its inhibitory effect on the TBK1-IRF3-STAT1 axis.

Notably, this regulatory axis is reciprocal. A recent preprint reports that PARP7 ADP-ribosylates p300 and CBP in the nucleus, promoting their ubiquitin-mediated turnover (Siordia *et al*., 2025). Together with our findings, these data define a bidirectional regulatory circuit in which p300/CBP-mediated acetylation stabilizes PARP7, thereby reinforcing repression of the TBK1-IRF3 pathway, while PARP7-mediated MARylation limits p300/CBP protein abundance and transcriptional output. Rather than representing a contradiction, this reciprocity suggests a substrate-dependent equilibrium whose functional outcome is shaped by cellular context. In tumor cells, where chronic cytosolic DNA stress and remodeled chromatin are prevalent, this circuit is likely biased toward persistent suppression of IFN-I signaling.

The broader biological context underscores the significance of this circuit. p300 and CBP are well-established co-activators of oncogenic transcription factors in cancer (Godfrey *et al*, 2025; Zhou *et al*, 2021). For example, they are essential co-activators for the androgen and estrogen receptors in prostate and breast cancers. More generally, p300/CBP are frequently overexpressed in multiple malignancies, where they drive proliferative, survival, metastatic and stress-response gene programs, and have been implicated in tumor immune evasion as well (Yuan *et al*, 2024). Classically, p300/CBP are known to co-operate with IRF3 and other factors to enhance IFN-β transcription during antiviral responses, but oncogenic reprogramming of the chromatin landscape can invert their role in tumors (Qin *et al*, 2005). Indeed, recent studies have begun to map p300/CBP-dependent enhancers for antigen presentation and cytokine genes, showing that p300/CBP activity can either promote or suppress anti-tumor immunity depending on context. Our results place PARP7 at the nexus of this immunological reprogramming. By stabilizing PARP7, p300/CBP help enforce persistent repression of the cGAS-STING-TBK1 pathway in cancer cells. Consistent with this model, pharmacological inhibition of p300/CBP’s HAT activity (for example by A-485) dramatically lowers PARP7 levels, unleashing IFN-β and CXCL10 induction in diverse tumor cell lines and markedly enhancing CD8^+^ T-cell infiltration and function in vivo.

We acknowledge several caveats. First, p300/CBP acetylate many substrates and broadly influence gene expression, so their effects on immunity likely extend beyond PARP7. Likewise, PARP7 itself ADP-ribosylates multiple transcriptional regulators, and whether acetylation alters its substrate specificity remains to be determined. Tumor-to-tumor variability in p300/CBP activity and in PARP7 expression also suggests that the net immune outcome of this pathway will be context-dependent. Nonetheless, several observations support our model. The half-life of PARP7 is exceptionally short (∼4.5 min), so even a modest increase in its acetylation (and thus stability) markedly raises the threshold for IFN induction (Yang *et al*., 2023).

Consistently, we find that p300/CBP inhibition robustly augments IFN-I signaling across tumor cell lines. These data argue that the p300/CBP–PARP7 axis is a key determinant of innate immune activation in cancer cells.

In summary, this work identifies a regulatory module in which p300/CBP-mediated acetylation stabilizes PARP7, and stabilized PARP7 dampens cGAS-STING-driven IFN-I signaling. When considered together with prior evidence that PARP7 can destabilize p300/CBP, our findings support a dynamic regulatory circuit in which reciprocal regulation between p300/CBP and PARP7 contributes to the maintenance of an immunosuppressive state in tumor cells. The net effect in tumor cells is a sustained suppression of IFN-I signaling. Therapeutically, this suggests new opportunities: targeting the p300/CBP-PARP7 node could reawaken antitumor immunity. In our study, pharmacological inhibition of p300/CBP HAT activity using A-485 was sufficient to activate IFN-I signaling in immunocompetent tumor models and to elicit measurable antitumor immune effects in vivo. Combining HAT inhibition with immune checkpoint blockade or STING agonists—or employing PARP7-selective inhibitors that disrupt this feedback loop —could yield synergistic antitumor effects, especially in immune “cold” tumors (Woo *et al*, 2014). Of course, given the broad roles of p300/CBP in normal and cancer cell biology, any such strategies will require careful testing in multiple tumor contexts. Nonetheless, our findings unveil a previously unrecognized regulatory circuit that tumors exploit to suppress innate immunity and point to the p300/CBP-PARP7 axis as a promising target to boost IFN-I-driven anticancer immunity.

## Acknowledgements

This work was supported by grants from the National Key Research and Development Program of China (2024YFC2815900) and the Scientific and Technological Research Program of Chongqing Municipal Education Commission (KJZD-M202400402). We thank Prof. Wei Yu and Prof. Shimin Zhao (Fudan University) for kindly providing the wild-type and mutant EP300 plasmids.

